# A thin line between conflict and reaction time effects on EEG and fMRI brain signals

**DOI:** 10.1101/2023.02.14.528515

**Authors:** Ewa Beldzik, Markus Ullsperger

## Abstract

The last two decades of electrophysiological and neuroimaging research converged that the activity in the medial frontal cortex plays a pivotal role in cognitive control processes. Notably, the midfrontal theta (MFT) oscillatory EEG power, as well as activity in the anterior midcingulate cortex (aMCC) or pre-supplementary motor area (preSMA), were consistently proclaimed as markers of conflict processing. However, these brain signals are strongly correlated with response time (RT) variability in various non-conflict tasks, which overshadows the true nature of their involvement. Our previous study (Beldzik et al., 2022) successfully identified these brain signals during a simultaneous EEG-fMRI experiment implementing Stroop and Simon tasks. Based on the assumption that overcoming the habitual prepotent response during high interference trials require additional neural resources beyond simple decision variable represented in RTs, here we aim to verify if these markers exhibit a congruency effect beyond RT variations. Furthermore, we explored if these brain signals represent either proactive or reactive cognitive control mechanisms by investigating two widely known behavioral phenomena observed in conflict tasks: proportion congruency and congruency sequence effects. The results revealed partially null findings for MFT activity, yet a distinct cognitive control specialization between aMCC and preSMA. Our study provides novel evidence that the former is involved in proactive control mechanisms, possibly contingency learning, whereas the latter reflects reactive control mechanisms by exhibiting a strong congruency effect regardless of RT variation and responding to adaptive behavior.

## 1. Introduction

In our daily life, we continuously rely on cognitive processes that allow us to adapt to changing environments and coordinate actions to optimize task goals. One of these vital processes is the human ability to detect and resolve conflict, which enables selecting relevant information despite a habitual tendency to another goal-irrelevant information. The last two decades of research in cognitive neuroscience cortex plays a pivotal role in these functions (for reviews, see Cespón et al., 2020; Heidlmayr et al., 2020). Many EEG and fMRI studies indicated that activity in this brain region increases for high-interference trials compared to low-interference trials. However, such comparison is confounded by reaction time (RTs) differences between these trial types. Considering that the medial frontal cortex shows a strong positive correlation with RT (Cohen and Cavanagh, 2011; Domagalik et al., 2014; Yarkoni et al., 2009), it is essential to investigate the conflict-related brain activity while accounting for the RT effect.

In a simplified term, a response conflict is a competition between mutually exclusive response options. In cognitive neuroscience, such competition can be evoked by commonly used conflict tasks, i.e., Stroop (Stroop, 1935), Flanker (Eriksen and Eriksen, 1974), or Simon (Simon, 1969). In these tasks, stimuli are presented with either a congruent feature (congruent trial) or a mismatched feature predisposing a competitive response (incongruent trials). The occurrence of the conflict is manifested by the prolonged reaction times (RTs) and higher error likelihood. Several brain markers were identified that resemble this characteristic as showing greater amplitude for incongruent trials in comparison to congruent ones. These include, but are not limited to, the N2 and N450 EEG potentials (Beldzik et al., 2015a; Folstein and Van Petten, 2008), midfrontal theta (MFT) oscillatory EEG power (Asanowicz et al., 2022, 2021; Cohen and Donner, 2013; Hanslmayr et al., 2008; Nigbur et al., 2011), as well activity in the anterior midcingulate cortex (aMCC) or pre-supplementary motor area (preSMA) as indicated by fMRI studies (Botvinick et al., 2004; Iannaccone et al., 2015; Nachev et al., 2008; Ullsperger et al., 2014; Ullsperger and Von Cramon, 2001). These *markers of conflict processing* were linked to various subfunctions in conflict monitoring, resolution, and subsequent adaptive behavior.

However, the increased activity for more difficult trials can be explained by sustained neural activity due to the ongoing neural computations and continuous inflow of metabolic supplies. For instance, neural response in the primary visual cortex to prolonged flashing checkboard will result in sustained visual gamma activity and linearly increasing amplitude of BOLD fMRI signal (Engell et al., 2012; Lewis et al., 2018; Logothetis et al., 2001). The same holds for responses of varying latency to stimuli of constant duration. The amplitude of the BOLD signal in many cortical regions shows a positive correlation to RTs across various cognitive tasks (Mumford et al., 2023; Yarkoni et al., 2009). This widespread positive RT-BOLD relationship was found even in the case of fast and homogenous saccadic responses (Domagalik et al., 2014). Finally, numerous M/EEG indicators present similar tendencies for changing linearly with decision time, e.g., centroparietal positivity (O’Connell et al., 2012; Twomey et al., 2015), non-phase locked MFT (Cohen and Donner, 2013; Duprez et al., 2020; Feuerriegel et al., 2021), or motor beta and gamma frequency bands (Donner et al., 2009; Fischer et al., 2018; Rogge et al., 2022).

Inspired by these proceedings, Grinband and colleagues (2011) verified whether conflict-related brain activations are not prone to this RT effect. The authors selected trials based on their RT scores and compared the two trial types when RTs were equalized or showed the opposite values of mean RTs. As a result, the aMCC activity equalized or was reversed, respectively, suggesting that this brain region is sensitive to the RT effect instead of being involved in conflict *per se*. Although this approach was questions by others (Yeung et al., 2011), similar conclusions were drawn by another fMRI study comparing the Multi-Source Interference task with a simple RT task (Carp et al., 2012; Weissman and Carp, 2013). The authors found that the RT-dependent increase of aMCC activity in the simple RT task fully accounts for the conflict-related aMCC increase in the interference task. Our research group investigated these effects for conflict-related activity evoked by a saccadic task (Beldzik et al., 2015b). Although we carefully followed the methods used in the study by Grinband et al. (2011), in contrast to their findings, all conflict-related brain activations, including preSMA, showed consistently greater activation for high interference stimuli. We concluded that preSMA reflects the pure congruency effect regardless of RT variations.

In a similar vein, a controversy was raised upon conflict-related MFT activity. Particularly, Scherbaum and Dshemuchadse (2013) simulated theta wavelets of different lengths, corresponding to longer RTs, to quantify the influence of RT on theta power. Although the amplitude of the theta signal was kept equal for all trials, incompatible trials showed greater energy than compatible ones. In response to these arguments, Cohen and Nigbur (2013) updated the model used by Scherbaum and Dshemuchadse in simulation and re-analyzed the criticized data from the Simon task, selecting trials with RTs from the same 50 ms time range. As a result, the conflict effect on MFT amplitude remained. Notably, previous studies using dipole fitting (Nigbur et al., 2011) or beamforming (Cohen and Ridderinkhof, 2013) tools have reported that the source of conflict-related MFT was estimated in the preSMA.

Thus, despite the substantial variability in methods employed, the abovementioned literature points toward a coherent picture: the aMCC activity is susceptible to RT effect, whereas the preSMA and MFT are not. To directly 1) verify the source of MFT activity and 2) test the coherency of that picture by evoking all conflict markers in a single experiment, we conducted a simultaneous EEG-fMRI study during conflict tasks while accounting for several conditions that varied in previous reports. The results addressing the first aim were recently described in Beldzik et al. (2022). Against the original hypothesis, yet in line with previous studies exploring the MFT-BOLD amplitude correlations, we only found a negative relationship of conflict-related MFT activity and activity in midline area 9, a brain region showing conflict-sensitive deactivation.

With the coherent picture disrupted, this study aimed to verify all conflict markers regarding the simple *RT effect* and two widely-known behavioral phenomena observed in conflict tasks: *proportion congruency* (Logan and Zbrodoff, 1979) and *congruency sequence* (Gratton et al., 1992; Ullsperger et al., 2005) *effects*. The proportion congruency effect is based on manipulating the frequency of congruent and incongruent trial occurrence to bias attention toward one of the stimulus features. As a result, RTs decrease for a more frequent type of trial and vice versa. Additionally, a list-wide proportion congruency effect is known to involve proactive global strategies operating before stimulus onset (Braver et al., 2007; Bugg et al., 2011, 2008). These strategies largely reflect expectations of the upcoming stimulus type. The congruency sequence effect, on the other hand, decreases interference on a trial if a high-interference trial precedes it. Such a situation triggers reactive control, i.e., short-term enhancement of the attentional set in reaction to conflict, which constitutes a fundamental mechanism for conflict adaptation (van den Wildenberg et al., 2012; Yang and Pourtois, 2022).

In our previous analysis with this dataset (Beldzik et al., 2022), we successfully identified MFT EEG activity and six independent fMRI brain networks sensitive to conflict. Here, we focused on three of those neural measures, the MFT amplitude, the aMCC, and preSMA networks’ activities, as they purport the brain markers of conflict processing commonly reported in the literature. Our goal was to verify if those markers exhibit 1) a congruency effect beyond RT variations and 2) proportion congruency and congruency sequence effects. The first goal was addressed in two fashions. First, the amplitude of each marker was compared after RT-based trial selection. We assumed that the *true* marker of conflict processing should have increased activity for high-interference trials despite similar RTs. Second, a linear mixed effect (LME) model was used to account for congruency and RT modulation in a timewise fashion. A conflict marker was expected to show greater LME coefficients for congruency than the RT condition. To address the study’s second goal, we run LME separately for proportion congruency and congruency sequence effects while controlling for RT variance in the model. We assumed that a conflict marker would exhibit either proactive or reactive cognitive control mechanism besides the primary congruency effect. Considering our previous study with oculomotor responses (Beldzik et al., 2015b), we hypothesized that preSMA is a marker that would validate these assumptions.

## 2. Methods

### 2.1 Participants

Thirty-seven participants (mean age, 22.1 ± 2.7 years; 22 females/15 male) met the following experiment requirements: no contraindication for MRI scanning, right-handedness (verified with the Edinburgh Handedness Inventory; Oldfield, 1971), normal or corrected-to-normal vision, no color-blindness (confirmed with Ishihara color vision test), no reported physical or psychiatric disorders, drug-free. Participants were informed about the procedure and goals of the study, and they gave written consent. The Bioethics Commission approved the study at Jagiellonian University. Data from two subjects were excluded during analysis due to the lack of frontocentral component in the EEG data (see Method section 2.5).

### 2.2 Experimental task

The task was prepared and generated using E-Prime 2.0 (Psychology Software Tools). It was presented on a 32-inch screen placed behind the MR scanner and approximately 100 cm from the head coil. Participants performed two types of conflict-inducing tasks, i.e., the Stroop (Fig. 1A) and the Simon (Fig. 1B) tasks. In the former, a stimulus was one of four color names (red, yellow, blue, or green) in Polish (Arial font, height 2°) printed in one of these colors. In the latter, a stimulus was a dot (diameter 2°) presented laterally (∼22°) to the fixation sign printed in one of these four colors. Although the tasks differed in stimulus features, they had the exact instructions given to participants: “Indicate an ink color of a stimulus ignoring its other features.” Indicating a color was obtained by pressing a specified button of response grip (Nordic Neuro Lab, Bergen, Norway) using a specified finger (left index finger, left thumb, right index finger, right thumb, respectively; Fig 1).

**Figure 1.**
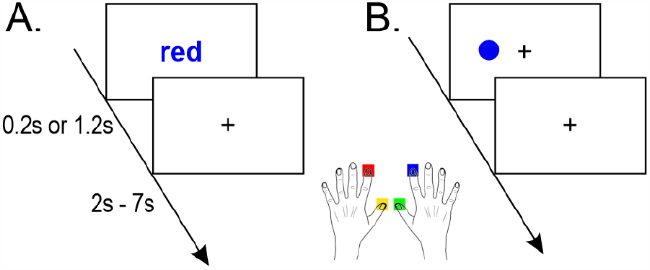
Scheme of the A) Stroop and B) Simon tasks.

Response conflict was present in trials where the target response was a semantic mismatch or contralateral to the stimulus location (“incongruent trials”). Conversely, no response conflict was present in trials where the target response was a semantic match or ipsilateral to the stimulus location (“congruent trials”). The stimulus was presented for either 200 ms (“short”) or 1200 ms (“long”). In both cases, the response window was 1.2 s. A ‘speed up’ icon was shown in case of a missing response. A fixation point (a plus sign, size 2° × 2°) was present throughout the whole experiment except for the stimuli presentation period in the Stroop task (see Fig. 1). The inter-trial time interval was randomly drawn from a uniform distribution of values in the following categories: 0.8, 1.3, 1.8, 2.3, 2.8, 5.3, and 5.8 s, resulting in an average of 3.5 s.

Each task was presented in two blocks of 50% and 20% congruency rates. Each session consisted of 60 incongruent and 60 or 240 congruent trials, respectively. These four sessions were counterbalanced between participants, with one restriction being the task type in the interleaved fashion (e.g., Stroop20%, Simon50%, Stroop50%, Simon20%). Between the sessions, subjects had an unlimited break from the task. Before beginning a new session, they were informed of the next stimulus type (word or dot) and a reminder with the response options. In total, the experiment lasted approximately 50 min and introduced 840 trials. Together, our design controlled for conflict (congruent vs. incongruent), task (Stroop vs. Simon), congruency rate (50% vs. 20%), and stimulus duration (short vs. long). The rationale for including these four conditions was to account for various task parameters, which differ in EEG and fMRI protocols.

Before the main experiment, participants undertook a training session, which included 12 centrally presented dots (neutral trials), 12 for the Simon task, and 12 for the Stroop task. If accuracy in each 12-trial run was below 90%, a subject had to redo the run.

### 2.3 EEG data acquisition and preprocessing

EEG data were recorded using an MR-compatible EEG cap (EasyCap, Herrsching, Germany) with 63 scalp electrodes, following the extended International 10-20 system and an additional channel for recording an electrocardiogram (ECG). The ECG electrode was placed on the participants’ back under the left shoulder blade to avoid signal contamination with chest movements. The reference electrode was positioned at FCz. EEG data were recorded using Vision Recorder (Version 1.20) with a sampling rate of 5 kHz. The electrode impedances were kept below 20 kΩ. The SyncBox (Brain Products GmbH, Gilching, Germany) was used to ensure that the EEG clock was synchronized to the MR scanner clock (Mullinger et al., 2008).

The first two steps of EEG data preprocessing were conducted using BrainVision Analyzer 2.0 software (Brain Products GmbH). MR artifacts were minimized using an average artifact subtraction (AAS) technique (Allen et al., 2000). Specifically, the gradient artifact was defined as a continuous interval of 1800 ms in length, beginning at the “start volume scan” marker. An artifact template was created using a sliding average of twenty-one artefactual intervals. Next, datasets were downsampled to 500 Hz. The AAS technique was also applied to correct ballistocardiogram artifacts. Again, twenty-one pulses in the semiautomatic mode were used to create a template. Peak detection was run on the ECG channel. EEG data were then exported to EEGLAB (version 2019.1; Delorme and Makeig, 2004), excluding the ECG channel. After removing resting periods, data were filtered in a 0.5 – 35 Hz range (*eegfiltnew*) and re-referenced by common average.

### 2.4 EEG component selection

The EEG data were temporarily epoched to mark ‘bad’ epochs. The criteria for a ‘bad’ epoch included either greater than three absolute normalized channel-mean variances in the period 500 ms before to 1200 ms after the stimulus onset (Nolan et al., 2010) or greater than 150 μV absolute amplitude in ± 100 ms relative to the response onset, accounting for possible movement-related artifacts (Beldzik et al., 2019). These criteria marked 8.0% (SD 8.1%) of the trials.

The ICA denoising approach was used to obtain a detectable frontocentral component for each participant (Beldzik et al., 2019; Scheeringa et al., 2016, 2011). First, continuous EEG data were bandpass filtered in the 4-8 Hz frequency range. Following epoch extraction in 0-1200 ms post-stimulus range and exclusion of ‘bad’ epochs and those with incorrect responses, the ICA was performed using the default, extended infomax algorithm (Lee et al., 1999). The unmixing weights were applied to the preprocessed data (i.e., before the denoising), and components were back-projected to the channel level. Applying the weights back on the original data enables time-frequency decomposition on the full power spectrum and extended epochs instead of theta-filtered short ones. The topographical maps were visually inspected for the most prominent frontocentral component. Out of 37 participants, two did not show any frontocentral components; thus, these subjects were removed from further analyses. Epochs were extracted from −1 s to 1.8 s relative to the stimulus presentation from the selected components’ time courses.

### 2.5 EEG time-frequency analysis

The time-frequency decomposition was carried out using the complex Morlet wavelet convolution. The frequency vector comprised 60 points in the 2-30 Hz range, increasing logarithmically. Similarly, the cycle values corresponding to each frequency ranged from 2-7, increasing logarithmically. The calculated spectral power was baseline-corrected by subtracting the mean power -500 to -200 ms before stimulus onset from each time point (Duprez et al., 2020). Next, the time-frequency plots were epoched from -800 to 400ms aligned to the response onset in 10ms resolution. The theta power was calculated as the mean amplitude power in the 4-8 Hz range for each time point, trial, and subject and underwent further analysis discussed in detail below. All analyses conducted here included correct trials only.

### 2.6 fMRI data acquisition

MRI was performed using a 3T scanner (Magnetom Skyra, Siemens) with a 64-channel head/neck coil. The isocenter was set 4 cm superior to the nasion to reduce gradient artifacts in the EEG data (Mullinger et al., 2011). High-resolution, whole-brain anatomical images were acquired using a T1-MPRAGE sequence. A total of 176 sagittal slices were obtained (voxel size 1 × 1 × 1.1 mm3; TR = 2,300 ms, TE = 2.98 ms, flip angle = 9°) for coregistration with the fMRI data. Next, a B0 inhomogeneity gradient field map (magnitude and phase images) was acquired with a dual-echo gradient-echo sequence, matched spatially with fMRI scans (TE1 = 4.92 ms, TE2 = 7.38 ms, TR = 508 ms).

Functional T2*-weighted images were acquired using a whole-brain echo-planar (EPI) pulse sequence with the following parameters: 3.5 mm isotropic voxel, TR = 1800ms, TE = 27 ms, flip angle = 75°, FOV 224 × 224 mm^2^, GRAPPA acceleration factor 2, and phase encoding A/P. Whole-brain images (cerebellum excluded) were covered with 34 axial slices taken in an interleaved order. Due to magnetic saturation effects, each session’s first three volumes (dummy scans) were instantly discarded, resulting in two sessions of 240 and two sessions of 590 volumes for each participant.

### 2.7 fMRI data preprocessing

Preprocessing of fMRI data was conducted using Analysis of Functional NeuroImage (AFNI, version 17.3.03; Cox, 1996) and the FMRIB Software Library (FSL, version 5.0.9; Jenkinson et al., 2012). Anatomical images were skull-stripped and coregistered to MNI (Montreal Neurological Institute) space using nonlinear transformation (*@SSwarper*). They were segmented (*FAST*) to create individual cerebrospinal fluid (CSF) masks. The first step of functional data preprocessing was to obtain the transformation matrix for motion correction (*3dvolreg*) to avoid its possible alteration by temporal interpolation applied further to fMRI data (Power et al., 2017). Next, de-spiking (*3dDespike*) and slice timing correction (*3dTshift*) were conducted. Then, transformation matrices for coregistration of functional data to anatomical data (*align_epi_anat*.*py*) as well as B0 inhomogeneity derived from gradient fieldmaps (*Fugue*) were calculated. The spatial transformation was performed in one step (*3dNwarpApply*), combining all prepared matrices, i.e., motion correction, anatomical co-registration, and distortion correction. The fMRI datasets were masked using a clip level fraction of 0.4, scaled to percent signal change, and the CSF signal was extracted using previously obtained individual masks. Finally, the functional images were coregistered to MNI space using the transformation matrix from nonlinear anatomical normalization.

To clear the fMRI signal from motion residuals, we applied ‘null’ regression (*3dREMLfit)* with the pre-whitening option (using ARMA_(1,1)_ model) to functional images. The model included twelve movement parameters (six demeaned originals and six first derivatives), the CSF time course, and four or nine polynomials as determined automatically using the ‘1 + int(D/150)’ equation, where D is the session’s duration. The rationale for regressing the CSF signal is that this signal reflects purely physiological noises, respiratory and cardiac, and often contains motion-related artifacts (Caballero-Gaudes and Reynolds, 2017; Power et al., 2014).

### 2.8 fMRI component selection

Our interim goal was to obtain a ‘functional parcellation’ of the BOLD signal in the frontal cortex to identify preSMA and aMCC regions in a data-driven fashion. Thus, a group ICA was conducted (GIFT version 4.0b; Calhoun et al., 2001) using a mask limited to all frontal, insular, and cingulate regions defined by the Harvard-Oxford cortical structural atlas (neurovault id: 1705). An estimation of the number of components was performed using minimum description length (MDL) criteria (Y.-O. Li et al., 2007). ICA decomposition stability was validated using ICASSO (Himberg et al., 2004) with 50 random initializations of the Infomax algorithm. Data were back-reconstructed using the default GICA option with z-scoring applied to both maps and time courses. The components’ maps were corrected with FDR at the α < .01 and inspected to identify and discard those primarily associated with artifacts representing signals from large vessels, ventricles, motion, and susceptibility (Griffanti et al., 2017; Kelly et al., 2010; Varoquaux et al., 2010). The time course of the brain’s components was interpolated to 100-ms resolution. Next, epochs were extracted from 0 s to 10 s of the stimulus onset and baseline corrected by subtracting the values at 0.

### 2.9 Trial selection analyses

The postprocessing of EEG and fMRI data comprised two parts. The first part focused on comparing the theta, aMCC, and preSMA amplitudes between congruent and incongruent trials while controlling for RT variance using the trial selection approach (Cohen and Nigbur, 2013; Grinband et al., 2011). First, a classical comparison between all congruent and incongruent trials was obtained for all three conflict markers. Epochs with theta power and hemodynamic responses were averaged for each condition and compared using a paired two-tailed t-test. The *p* values corresponding to each time-point were corrected with FDR at the α < .05. Second, a similar analysis was conducted, only here RTs for each trial type were equalized. Selection of trials was performed by normalizing RTs for each participant and including congruent trials within -0.6 – 1.3 z-values and incongruent trials within -1.5 – 1.4 z-values. These values were estimated to ensure a nonsignificant RT difference between the congruency conditions and a maximal number of trials for the comparison (58% of congruent and 78% of incongruent trials remained after selection). Third, data were separated into quantiles based on the normalized RT values of incongruent trials, and congruent trials were selected to match RT values in each bin. This last comparison was conducted only for theta and BOLD peak amplitudes to compensate for the lost statistical power due to trial selection.

### 2.10 LME analyses

The second part of EEG and fMRI data analyses was applied using a linear mixed-effect (LME) model to investigate neural processes related to conflict adaptation on a trial-by-trial level. Particularly, theta, aMCC, and preSMA amplitudes underwent the LME model to account for 1) RT variability, 2) proportion congruency effect, and 3) congruency sequence effect. The model is ideal for capturing small effect sizes as it enables the inclusion of single-trial measurements in one group analysis. It has proven useful in previous studies (Beldzik et al., 2022, 2019). The RT model was a simple formula applied to all three brain measures at each time point within their epochs:

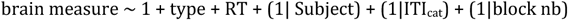

Since RT values were normalized, congruency and RTs estimates could be compared to make inferences about the dominant effect. We assumed that if the estimates for conflict were higher than the estimates of RT, it would speak in favor of the marker of conflict. Otherwise, this brain marker is primarily representing the time of neural computations or could simply be confounded by RTs due to spurious correlations with the hemodynamic response model (Mumford et al., 2023).

Next, we examined if these brain measures showed more sophisticated conflict-related effects, that is, proportion congruency and congruency sequence effects. To maximize their sensitivity, we marked the time point with maximal congruency effect for each brain measure and extracted the values in the 100 ms time range around it. Such values underwent LME models with the following formulas:

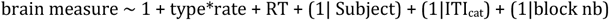

for the proportion congruency effect, and

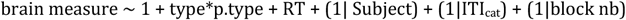

for the congruency sequence effect, where *p*.*type* denotes the previous trial type. Additionally, RT values underwent similar LME analyses. The final matrix comprised 24322 observations.

## 3. Results

### 3.1 Defining conflict markers of interest

A comprehensive description of the results of the EEG and fMRI data analysis was presented in our previous work (Beldzik et a, 2022). Here we focus only on three conflict markers identified before and widely referenced in the literature, i.e., 1) response-locked theta power, 2) aMCC, and 3) preSMA hemodynamic responses. The selected EEG components with frontocentral topography showed a maximum at the FCz channel and pronounced activity in the response-locked theta power (Fig. 2A). The fMRI results revealed two distinct components involving the cortical loci of interest (Table 1; Fig. 2B). The aMCC component covered a single region, whereas the preSMA was functionally coupled to the bilateral ventrolateral prefrontal cortices. For simplicity, we shall refer to this component as the preSMA network. Notably, in the study by Sallet and colleagues (2013), the authors investigated the resting-state connectivity of all subregions within the medial frontal wall. They found that the preSMA was coupled to the caudal ventral prefrontal cortex centered at [49, 31, 19]. These coordinates are in close proximity to the coordinates of the ventrolateral prefrontal cortex found here [49, 31, 22].

**Table 1.** Brain regions corresponding to fMRI brain components.

| Label | Region | Side | x | y | z | t |
| --- | --- | --- | --- | --- | --- | --- |
| aMCC | cingulate gyrus | M | 3 | 24 | 33 | 16.7 |
| preSMA | pre-supplementary motor area | M | -4 | 24 | 50 | 15.7 |
|  | inferior frontal sulcus | L | -49 | -10 | 33 | 16.1 |
|  |  | R | 49 | 31 | 22 | 13.3 |

Note. R=right; L=left; M=medial; A = area; x.y.z coordinates are provided in MNI space. preSMA=pre-supplementary motor area. aMCC=anterior midcingulate cortex.

**Figure 2.**
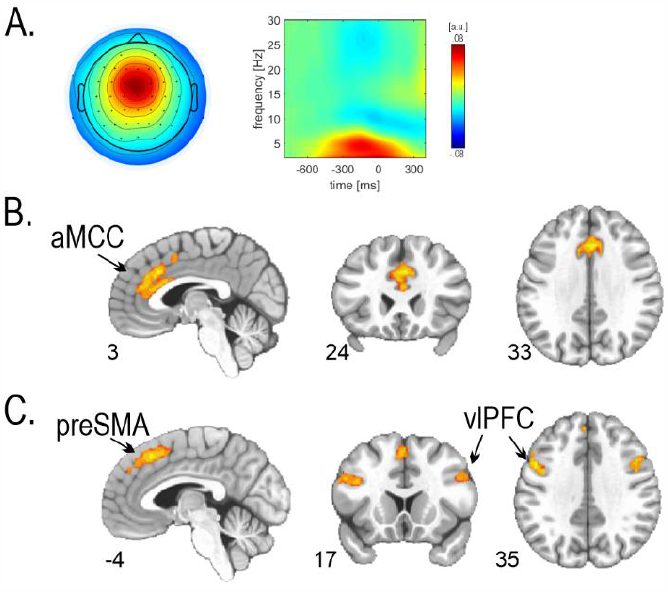
Conflict markers of interest. A) Midfrontal EEG component topography (left) and response-locked spectral power (right). fMRI components corresponding to B) aMCC and C) preSMA. preSMA=pre-supplementary motor area. aMCC=anterior midcingulate cortex. vlPFC=ventrolateral prefrontal cortex.

### 3.2 Trial selection results

Participants (N=35) committed 5.9% (SD 3.9%) erroneous responses and 3.9% (SD 3.2%) omissions. All results reported here are based on correct trials only. In line with the assumption of conflict processing, congruent and incongruent trials differed substantially in their mean RTs (congruent: 641.0; SD 59.0 ms; incongruent 728.4, SD 57.4 ms ; t(34) = 18.8, p < .001; d_cohen_=3.2). For this comparison, all three brain measures showed significantly greater activity for incongruent trials than for congruent ones (Fig. 3A). In the next comparison, we selected trials to equalize their group-mean RTs (congruent: 673.6 ms; SD 62.1 ms; incongruent 673.8 ms; SD 60.6 ms; t=0.1, p=.92; d_cohen_=0.02). As a result, conflict-related differences in theta and preSMA activity weakened yet remained significant (Fig. 3B). In contrast, the difference between congruent and incongruent trials in the case of aMCC activity vanished at the peak of hemodynamic response but remained significant at its undershoot. The third comparison was conducted between quantile bins based on the normalized RTs. The results revealed that the only brain measure which showed persistent conflict-related sensitivity at each bin was the preSMA (Fig. 3C). The midfrontal theta, on the other hand, showed a profound congruency effect but only in the case of the fastest trials.

**Figure 3.**
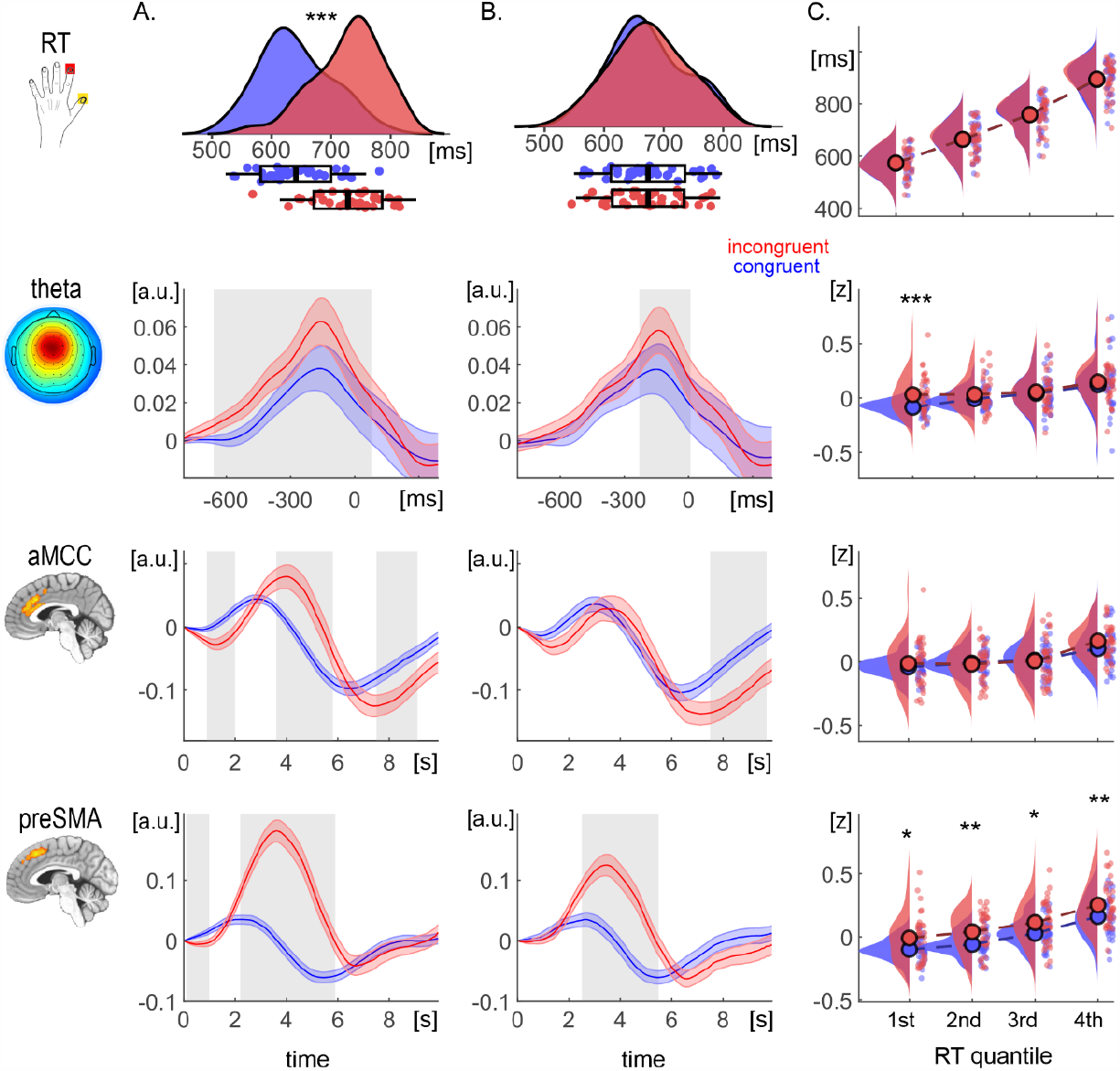
Results of the trial selection analyses aimed to verify the RT effect. Congruency effect for the amplitude of the MFT, aMCC, and preSMA in the case for A) all trials, B) trials with equalized RT, and C) quantiles with equalized RT (peak activity only). The top row presents the corresponding RT values. Bars denote standard errors. The shaded areas represent FDR-corrected significant t-tests (p_cor_ <.05). *** p <.001; ** p <.005; * p <.05. MFT=midfrontal theta; aMCC= anterior Midcingulate Cortex; preSMA= pre-supplementary motor area.

### 3.3 LME results

In the second part of data postprocessing, we conducted data analyses with three approaches using the LME model. The first LME analysis investigating congruency and RT effects at each time point of brain amplitudes revealed a general temporal overlap of the two effects (Fig 4A). Interestingly, only the theta and preSMA exhibited greater estimate values for the congruency effect than for the RT effect. Further LME analyses were conducted only for the amplitude values at the peaks of congruency effects. Specifically, the maximal congruency effects were observed at -135 ms before the response in theta activity and 4.5 s and 4.0 s after the stimulus in aMCC and preSMA, respectively.

**Figure 4.**
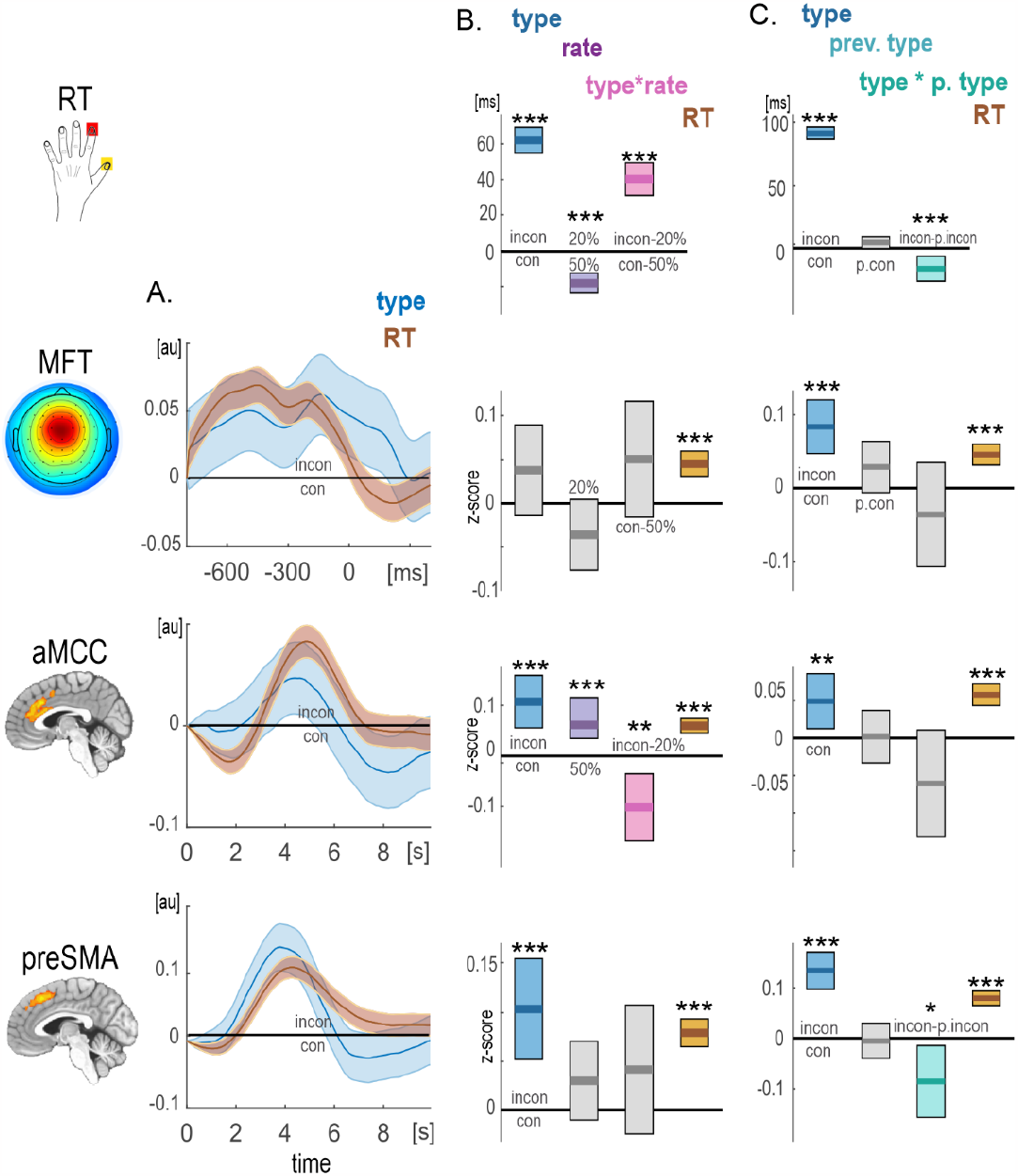
Results of the LME analyses aimed to verify the RT, proportion congruency, and congruency sequence effects. A) Estimates for the trial type and normalized reaction times (RTs) effects predicting the amplitude of the MFT, aMCC, and preSMA) in a point-by-point fashion. B) Estimates for the raw RTs (1^st^ row) and the amplitude of each brain measure (2^nd^ - 4^th^ rows) at the peak of congruency effects exploring B) the proportion congruency and C) congruency sequence effects. Bars denote confidence intervals. *** p <.001; ** p <.005; * p <.05. MFT=midfrontal theta; aMCC= anterior Midcingulate Cortex; preSMA= pre-supplementary motor area.

The second LME analysis investigating the proportion congruency effect was applied to RT scores and three brain measures (Table 2; Fig 4B). In general, responses were considerably faster in blocks with a 20% congruency rate than in 50% blocks indicating the facilitation process related to the sequence of congruent trials. Most importantly and in line with the proportion congruency effect, responses for incongruent trials were slower (in addition to the general conflict effect) in 20% blocks than congruent trials in 50% blocks, as indicated by the interaction term. This finding can be linked with increased interference for 20% of blocks. The MFT activity showed the general decrease in 20% blocks compared to the 50% blocks. Interestingly, the aMCC activity exhibited a pronounced increase in high-congruent block in general and a decrease for high-interference trials in that block as compared to low-congruent block and low-interference trials, respectively.

**Table 2.** Results of LME analysis investigating the proportion congruency effect.

| Dependent | Predictor | Estim | SE | t | DF | pValue | Lower | Upper |
| --- | --- | --- | --- | --- | --- | --- | --- | --- |
| RT (ms) | type_incon | 62.14 | 3.64 | 17.07 | 20247 | .001 | 55.0 | 69.3 |
|  | rate_20% | -17.78 | 2.86 | -6.22 | 20247 | .001 | -23.4 | -12.2 |
|  | type_incon:rate_20% | 40.08 | 4.68 | 8.56 | 20247 | .001 | 30.9 | 49.3 |
| MFT | RT | 0.05 | 0.01 | 6.19 | 20245 | .001 | 0.03 | 0.06 |
| aMCC | type_incon | 0.11 | 0.03 | 4.05 | 20245 | .001 | 0.06 | 0.16 |
|  | rate_20% | 0.08 | 0.02 | 3.63 | 20245 | .001 | 0.03 | 0.12 |
|  | type_incon:rate_20% | -0.10 | 0.03 | -3.00 | 20245 | .003 | -0.17 | -0.04 |
|  | RT | 0.06 | 0.01 | 8.14 | 20245 | .001 | 0.05 | 0.07 |
| preSMA | type_incon | 0.10 | 0.03 | 3.93 | 20245 | .001 | 0.05 | 0.16 |
|  | RT | 0.08 | 0.01 | 10.89 | 20245 | .001 | 0.07 | 0.09 |

Note. Only significant estimates are presented. Estim = parameter estimate. SE = standard error. DF = degrees of freedom. Lower/Upper = lower/upper confidence interval. MFT=midfrontal theta; aMCC=anterior midcingulate cortex; preSMA=pre-supplementary motor area

The final LME analysis investigating the congruency sequence effect was applied to brain measures and raw RT scores (Table 3; Fig. 4C). In line with the congruency sequence effect, when preceded by an incongruent trial, responses to incongruent stimuli speeded profoundly. Also, there was a marginal post-conflict slowing of congruent trials. These results suggest behavioral adaptation and loss of interference following incongruent trials. Regarding the brain measures, only the activity in preSMA showed this interaction effect as significant (Fig. 4C, bottom row). Notably, all LME results reported here show similar results even when normalized RTs are included in the model.

**Table 3.** Results of LME analysis investigating the congruency sequence effect.

| Dependent | Predictor | Estim | SE | t | DF | pValue | Lower | Upper |
| --- | --- | --- | --- | --- | --- | --- | --- | --- |
| RT (ms) | type_incon | 91.37 | 2.51 | 36.36 | 20247 | .001 | 86.4 | 96.3 |
|  | type_incon:rate_20% | -16.14 | 5.04 | -3.20 | 20247 | .001 | -26.0 | -6.3 |
| MFT | type_incon | 0.08 | 0.02 | 4.48 | 20245 | .001 | 0.05 | 0.12 |
|  | RT | 0.05 | 0.01 | 6.26 | 20245 | .001 | 0.03 | 0.06 |
| aMCC | type_incon | 0.05 | 0.02 | 2.63 | 20245 | .009 | 0.01 | 0.09 |
|  | RT | 0.06 | 0.01 | 7.91 | 20245 | .001 | 0.04 | 0.07 |
| preSMA | type_incon | 0.13 | 0.02 | 7.19 | 20245 | .001 | 0.10 | 0.17 |
|  | type_incon:rate_20% | -0.09 | 0.04 | -2.36 | 20245 | .018 | -0.16 | -0.01 |
|  | RT | 0.08 | 0.01 | 10.91 | 20245 | .001 | 0.07 | 0.09 |

Note. Only significant estimates are presented. Estim = parameter estimate. SE = standard error. DF = degrees of freedom. Lower/Upper = lower/upper confidence interval. MFT=midfrontal theta; aMCC=anterior midcingulate cortex; preSMA=pre-supplementary motor area

## 4. Discussion

The results presented here are a follow-up of previously described work (Beldzik et al., 2022). That study was based on the same dataset and focused on coupling MFT activity with activity in all brain regions or networks identified using fMRI in the frontal, cingulate, or insular cortices. Against previous claims based on MFT source estimations in the preSMA or aMCC, no such relationship was observed in either of these brain regions. Instead, we found a consistent negative correlation of conflict MFT with the BOLD amplitude in the midline area 9, a brain region showing conflict-sensitive deactivation and omission-preceding activation. Such a relationship has been reported in previous simultaneous EEG-fMRI studies. Specifically, MFT amplitude showed negative correlations to BOLD signal in the default mode network, including the midline prefrontal cortex, during the resting state (Prestel et al., 2018; Scheeringa et al., 2008), working memory (Scheeringa et al., 2009), and decision (Algermissen et al., 2021) tasks. Scheeringa and colleagues (2009) suggested that theta serves a role in task-orientedness by inhibiting irrelevant information and enabling optimal performance on a task. Indeed, midline area 9 is considered a subsystem of the default mode network (Andrews-Hanna et al., 2014) that was linked to mind-wandering (Durantin et al., 2015; Li et al., 2007). Thus, we concluded that the negative relationship of this brain area with conflict theta suggests that MFT plays a role in an active inhibition of the self-referential and mind-wandering processes that may distract participants from the task at hand and lead to omission error (Beldzik et al., 2022).

The results obtained here regarding RT, proportion congruency, and congruency sequence effects on MFT activity are not against that conclusion. In line with Nigbur and Cohen (2013), the MFT showed increased activity for incongruent trials in comparison to congruent ones for trials with equalized RTs (Fig. 3B). However, more detailed analysis with multiple RT bins indicated the difference only for the case of the fastest trials (Fig. 3C). Next, the LME analysis revealed that congruency and RT factors contribute similarly to the MFT variance (Fig. 4A). The following LME analyses showed a lack of MFT findings regarding both effects of interest (Fig. 4B, C). The null congruency sequence effect is in opposition to previous EEG studies (Gyurkovics and Levita. 2021; van Driel et al., 2015), and it may be a result of the generally low quality of EEG data recorded in the magnetic field environment. Nevertheless, the MFT results obtained here provide little evidence for theta reflecting a conflict-related process. Instead, as suggested by our previous study (Beldzik et al., 2022), MFT may be involved in the active inhibition of self-reflective cognition that may otherwise disrupt optimal performance. An alternative recently proposed interpretation for theta oscillations is that it reflects passive cortical disengagement, where an entire network or brain area ceases to receive inputs and essentially goes in “standby,” similar to occipital alpha waves (Snipes et al., 2022). Thus, MFT may comprise a very specific control signal rather than a conflict signal *per se*.

The aMCC brain activity showed consistent sensitivity to the RT effect. In line with Grinband and colleagues (2011), the conflict effect disappeared when aMCC activity was compared to trials with equalized RTs (Fig. 3B. C). Similarly, the LME model indicated that the RT factor preponderated the congruency effect (Fig. 4A). Interestingly, aMCC showed a prominent yet opposite direction to RTs scores, proportion congruency effect, and no congruency sequence effect. This former finding fits well with the theoretically and behaviorally outlined accounts of proactive control mechanisms in conflict, particularly contingency learning (Braem et al., 2019; Bugg, 2017; Bugg and Crump, 2012; Schmidt, 2013). The aMCC seems to play a role in anticipating when the next incongruent stimulus would appear, the process that is enhanced for blocks with a trial frequency ratio above the chance level and decreased for anticipated trials, that is, for incongruent trials in 20% block. This finding fits well with the idea that aMCC represents the value of expectancy violation (Vassena et al., 2020), i.e., when incongruent trials are expected and a congruent one comes, this is surprising and may trigger some adjustments.

Finally, the preSMA and bilateral inferior frontal sulci activity showed a robust congruency sequence effect despite fully accounting for RT variance in the data. In line with our previous study using oculomotor responses (Beldzik et al., 2015b), preSMA increased for incongruent trials in comparison to congruent for all RT bins (Fig. 3C). Also, the congruency effect dominated over RT effect in LME model (Fig. 4A). This has important implications in a debate over the ‘concept of conflict’ introduced by Grinband and colleagues (2011). The fact that RT variance could explain the conflict-related activity in aMCC activity had posed a serious challenge to the conflict-monitoring hypothesis (Botvinick et al., 2004). Yeung, Cohen & Botvinick (2011) replied that conflict tracks RTs variance more closely than the stimulus categorization. The authors declared that when simulated appropriately, the conflict monitoring theory predicts precisely the patterns of results presented by Grinband and colleagues (2011). In reply to their response, Grinband and co-workers (2011b) pointed out that similar activity can be observed in simple response task, which do not introduce any conflict. Moreover, the authors concluded that defining conflict as “any sensorimotor or cognitive process that lengthens RT” trivializes the idea of conflict and weakens its usefulness as a psychological construct. With such definition, it would be impossible to differentiate conflict from any other sensorimotor, memory, or attentional processes.

Here we evoked preSMA conflict-related activity that is independent of RT variance. It proves that conflict is a process that engages additional neural resources to overcome the biased response even when RTs were not lengthened. Moreover, we found that preSMA showed decreased activity in the incongruent trial that followed another incongruent trial. The congruency sequence effect was observed even when controlling for RT variance (Fig 4C). The congruency sequence effect is typically thought to measure a short-lived, reactive adaptation to a just-experienced conflict between competing response representations (Braem et al., 2019; van den Wildenberg et al., 2012; Yang and Pourtois, 2022). Thus, our findings indicated that preSMA is involved in conflict adaptation. Notably, in a recent study by Fu and colleagues (2022), the spiking of neuronal assemblies was directly recorded in the medial frontal cortex of epilepsy patients performing a Stroop task. Although the authors found that both aMCC and preSMA areas independently supported compositional conflict coding, the former demonstrated pre-response conflict information first, whereas the latter exhibited post-conflict information first. Their results obtained here align well with the proactive and reactive functional distinction of aMCC and preSMA brain regions.

## 5. Conclusions

The study verified three highly replicable conflict markers for congruency, proportion congruency, and congruency sequence effects while controlling for RT variance. The pre-response MFT EEG activity showed a dominance of the RT effect in its variance, except for the fastest response bin. The fMRI data revealed a distinct cognitive control specialization between the two brain regions of interest. The aMCC activity was increased for blocks where congruent trials occurred five times more often than incongruent ones. Although its activity showed a general increase for high-interference trials compared to low-interference, this effect was attenuated in 20% congruency ratio blocks. In contrast, preSMA manifested a significant congruency sequence effect even though RTs were included in the model. Together, our results indicate that aMCC is involved in proactive expectation of rare stimuli, whereas preSMA is responsible for reactive control by resolving the conflict and adapting to it both on the behavioral and neural levels.

## Data availability

The source data are also publicly available at https://osf.io/cx8a9/

## Code availability

All code generated for this study’s analyses are publicly available at https://github.com/ewabeldzik/conflict_RT_effects

## Acknowledgments

This project has received funding from the National Science Centre Poland (grant 2016/21/D/HS6/02962, to EB) and from the European Research Council (ERC) under the European Union’s Horizon 2020 research and innovation programme (grant agreement No 101018805, to MU). We thank Anna Bereś, Laura Łępa, and Magdalena Wielgus for their assistance with data collection. The authors declare no conflicts of interest related to this manuscript.

---

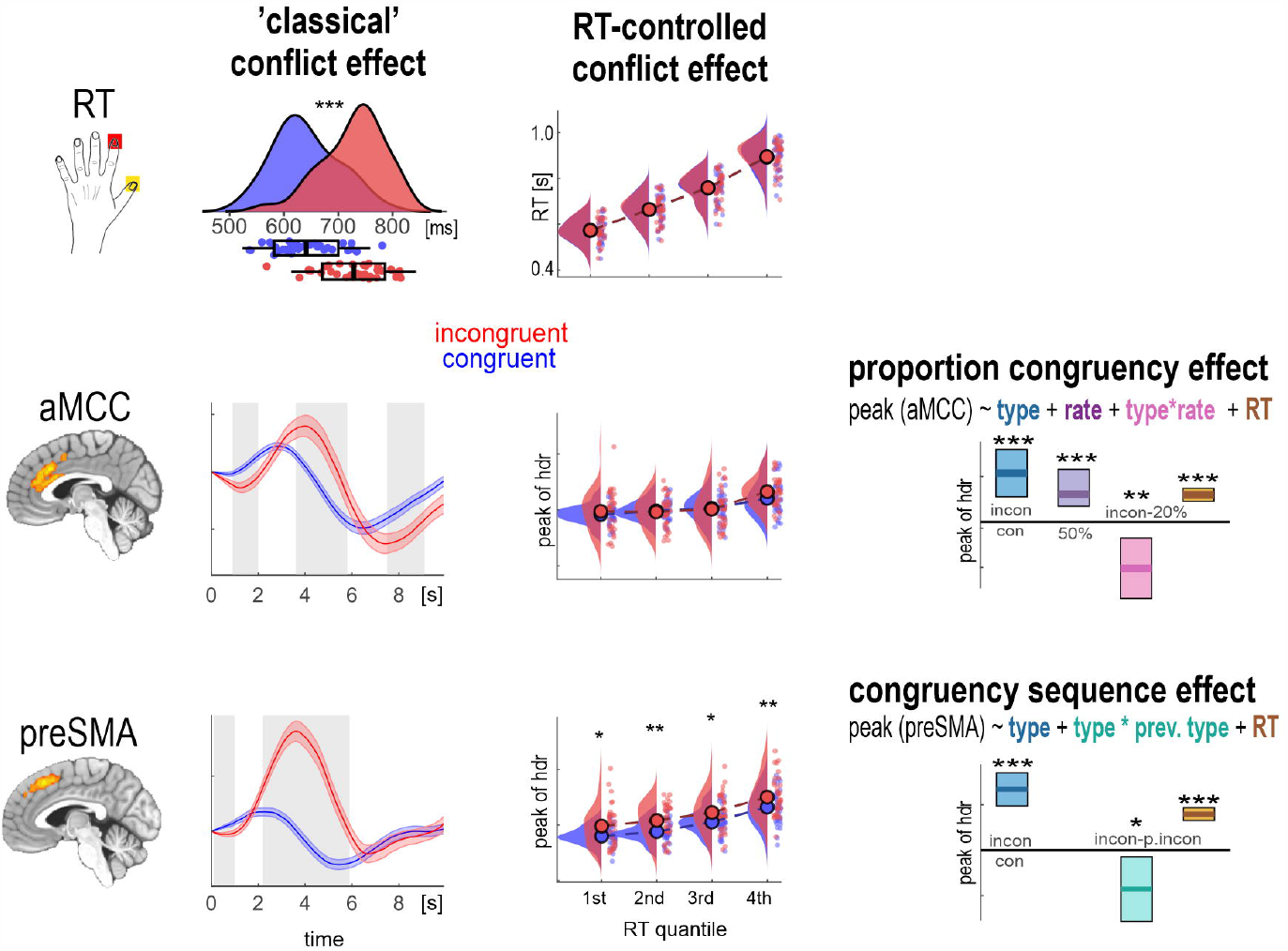

## Notes

### Competing Interest Statement

The authors have declared no competing interest.

### Summary of Updates

The following very relevant citations have been added to the manuscript: - Mumford et al., 2023 - Snipes et al., 2022 - Asanowicz et al., 2021; 2022 - Feuerriegel et al., 2021

## References

Algermissen, J., Swart, J.C., Scheeringa, R., Cools, R., Ouden, H.E.M. den, 2021. Striatal BOLD and midfrontal theta power express motivation for action. bioRxiv 2020.09.11.292870. https://doi.org/10.1101/2020.09.11.292870

Allen, P.J., Josephs, O., Turner, R., 2000. A Method for Removing Imaging Artifact from Continuous EEG Recorded during Functional MRI. Neuroimage 12, 230–239. https://doi.org/10.1006/NIMG.2000.0599

Andrews-Hanna, J.R., Smallwood, J., Spreng, R.N., 2014. The default network and self-generated thought: Component processes, dynamic control, and clinical relevance. Ann. N. Y. Acad. Sci. 1316, 29–52. https://doi.org/10.1111/nyas.12360

Asanowicz, D., Kotlewska, I., Panek, B., 2022. Neural underpinnings of proactive and preemptive adjustments of action control. J. Cogn. Neurosci. 34, 1590–1615. https://doi.org/10.1162/jocn_a_01884

Asanowicz, D., Panek, B., Kotlewska, I., 2021. Selection for action: The medial frontal cortex is an executive hub for stimulus and response selection. J. Cogn. Neurosci. 33, 1442–1469. https://doi.org/10.1162/jocn_a_01727

Beldzik, E., Domagalik, A., Beres, A., Marek, T., 2019. Linking visual gamma to task-related brain networks—a simultaneous EEG-fMRI study. Psychophysiology 56. https://doi.org/10.1111/psyp.13462

Beldzik, E., Domagalik, A., Froncisz, W., Marek, T., 2015a. Dissociating EEG sources linked to stimulus and response evaluation in numerical Stroop task using Independent Component Analysis. Clin. Neurophysiol. 126, 914–26. https://doi.org/10.1016/j.clinph.2014.08.009

Beldzik, E., Domagalik, A., Oginska, H., Marek, T., Fafrowicz, M., 2015b. Brain activations related to saccadic response conflict are not sensitive to time on task. Front. Hum. Neurosci. 9, 664. https://doi.org/10.3389/fnhum.2015.00664

Beldzik, E., Ullsperger, M., Domagalik, A., Marek, T., 2022. Conflict- and error-related theta activities are coupled to BOLD signals in different brain regions. Neuroimage 256, 119264. https://doi.org/10.1016/j.neuroimage.2022.119264

Botvinick, M.M., Cohen, J.D., Carter, C.S., 2004. Conflict monitoring and anterior cingulate cortex: an update. Trends Cogn. Sci. 8, 539–46.

Braem, S., Bugg, J.M., Schmidt, J.R., Crump, M.J.C., Weissman, D.H., Notebaert, W., Egner, T., 2019. Measuring Adaptive Control in Conflict Tasks. Trends Cogn. Sci. https://doi.org/10.1016/j.tics.2019.07.002

Braver, T.S., Gray, J.R., Burgess, G.C., 2007. Explaining the Many Varieties of Working Memory Variation: Dual Mechanisms of Cognitive Control, in: Conway, A.R.A., Jarrold, C., Kane, M.J., Miyake, A., Towse, J.N. (Eds.), Variation in Working Memory. Oxford University Press, pp. 76–106.

Bugg, J.M., 2017. Context, Conflict, and Control, in: The Wiley Handbook of Cognitive Control. John Wiley & Sons, Ltd, pp. 79–96. https://doi.org/10.1002/9781118920497.ch5

Bugg, J.M., Crump, M.J.C., 2012. In Support of a Distinction between Voluntary and Stimulus-Driven Control: A Review of the Literature on Proportion Congruent Effects. Front. Psychol. 3, 367. https://doi.org/10.3389/fpsyg.2012.00367

Bugg, J.M., Jacoby, L.L., Chanani, S., 2011. Why it is too early to lose control in accounts of item-specific proportion congruency effects. J. Exp. Psychol. Hum. Percept. Perform. 37, 844–859. https://doi.org/10.1037/a0019957

Bugg, J.M., Jacoby, L.L., Toth, J.P., 2008. Multiple levels of control in the Stroop task. Mem. Cogn. 36, 1484–1494. https://doi.org/10.3758/MC.36.8.1484

Caballero-Gaudes, C., Reynolds, R.C., 2017. Methods for cleaning the BOLD fMRI signal. Neuroimage 154, 128–149. https://doi.org/10.1016/j.neuroimage.2016.12.018

Calhoun, V.D., Adali, T., Pearlson, G.D., Pekar, J.J., 2001. Spatial and temporal independent component analysis of functional MRI data containing a pair of task-related waveforms. Hum. Brain Mapp. p13, 43–53. https://doi.org/10.1002/hbm.1024

Carp, J., Fitzgerald, K.D., Taylor, S.F., Weissman, D.H., 2012. Removing the effect of response time on brain activity reveals developmental differences in conflict processing in the posterior medial prefrontal cortex. Neuroimage 59, 853–60.

Cespón, J., Hommel, B., Korsch, M., Galashan, D., 2020. The neurocognitive underpinnings of the Simon effect: An integrative review of current research. Cogn. Affect. Behav. Neurosci. https://doi.org/10.3758/s13415-020-00836-y

Cohen, M.X., Cavanagh, J.F., 2011. Single-Trial Regression Elucidates the Role of Prefrontal Theta Oscillations in Response Conflict. Front. Psychol. 2, 30. https://doi.org/10.3389/fpsyg.2011.00030

Cohen, M.X., Donner, T.H., 2013. Midfrontal conflict-related theta-band power reflects neural oscillations that predict behavior. J. Neurophysiol. 110, 2752–2763. https://doi.org/10.1152/jn.00479.2013

Cohen, M.X., Nigbur, R., 2013. Reply to “Higher response time increases theta energy, conflict increases response time.” Clin. Neurophysiol. https://doi.org/10.1016/j.clinph.2013.03.013

Cohen, M.X., Ridderinkhof, K.R., 2013. EEG Source Reconstruction Reveals Frontal-Parietal Dynamics of Spatial Conflict Processing. PLoS One 8, e57293. https://doi.org/10.1371/journal.pone.0057293

Cox, R.W., 1996. AFNI: software for analysis and visualization of functional magnetic resonance neuroimages. Comput. Biomed. Res. 29, 162–173.

Delorme, A., Makeig, S., 2004. EEGLAB: an open source toolbox for analysis of single-trial EEG dynamics including independent component analysis. J. Neurosci. Methods 134, 9–21. https://doi.org/10.1016/j.jneumeth.2003.10.009

Domagalik, A., Beldzik, E., Oginska, H., Marek, T., Fafrowicz, M., 2014. Inconvenient correlation - RT-BOLD relationship for homogeneous and fast reactions. Neuroscience 278. https://doi.org/10.1016/j.neuroscience.2014.08.012

Donner, T.H., Siegel, M., Fries, P., Engel, A.K., 2009. Buildup of Choice-Predictive Activity in Human Motor Cortex during Perceptual Decision Making. Curr. Biol. 19, 1581–1585. https://doi.org/10.1016/j.cub.2009.07.066

Duprez, J., Gulbinaite, R., Cohen, M.X., 2020. Midfrontal theta phase coordinates behaviorally relevant brain computations during cognitive control. Neuroimage 207, 116340. https://doi.org/10.1016/j.neuroimage.2019.116340

Durantin, G., Dehais, F., Delorme, A., 2015. Characterization of mind wandering using fNIRS. Front. Syst. Neurosci. 9, 45. https://doi.org/10.3389/fnsys.2015.00045

Engell, A.D., Huettel, S., McCarthy, G., 2012. The fMRI BOLD signal tracks electrophysiological spectral perturbations, not event-related potentials. Neuroimage 59, 2600–6. https://doi.org/10.1016/j.neuroimage.2011.08.079

Eriksen, B.A., Eriksen, C.W., 1974. Effects of noise letters upon the identification of a target letter in a nonsearch task. Percept. Psychophys. 16, 143–149. https://doi.org/10.3758/BF03203267

Feuerriegel, D., Jiwa, M., Turner, W.F., Andrejević, M., Hester, R., Bode, S., 2021. Tracking dynamic adjustments to decision making and performance monitoring processes in conflict tasks. Neuroimage 238, 118265. https://doi.org/10.1016/j.neuroimage.2021.118265

Fischer, A.G., Nigbur, R., Klein, T.A., Danielmeier, C., Ullsperger, M., 2018. Cortical beta power reflects decision dynamics and uncovers multiple facets of post-error adaptation. Nat. Commun. 9, 5038. https://doi.org/10.1038/s41467-018-07456-8

Folstein, J.R., Van Petten, C., 2008. Influence of cognitive control and mismatch on the N2 component of the ERP: a review. Psychophysiology 45, 152–70.

Fu, Z., Beam, D., Chung, J.M., Reed, C.M., Mamelak, A.N., Adolphs, R., Rutishauser, U., 2022. The geometry of domain-general performance monitoring in the human medial frontal cortex. Science (80-.). 376. https://doi.org/10.1126/science.abm9922

Gratton, G., Coles, M.G., Donchin, E., 1992. Optimizing the use of information: Strategic control of activation of responses. J. Exp. Psychol. Gen. 121, 480–506. https://doi.org/10.1037//0096-3445.121.4.480

Griffanti, L., Douaud, G., Bijsterbosch, J., Evangelisti, S., Alfaro-Almagro, F., Glasser, M.F., Duff, E.P., Fitzgibbon, S., Westphal, R., Carone, D., Beckmann, C.F., Smith, S.M., 2017. Hand classification of fMRI ICA noise components. Neuroimage 154, 188–205. https://doi.org/10.1016/J.NEUROIMAGE.2016.12.036

Grinband, J., Savitskaya, J., Wager, T.D., Teichert, T., Ferrera, V.P., Hirsch, J., 2011. The dorsal medial frontal cortex is sensitive to time on task, not response conflict or error likelihood. Neuroimage 57, 303–11. https://doi.org/10.1016/j.neuroimage.2010.12.027

Gyurkovics, M., Levita, L., 2021. Dynamic Adjustments of Midfrontal Control Signals in Adults and Adolescents. Cereb. Cortex 31, 795–808. https://doi.org/10.1093/cercor/bhaa258

Hanslmayr, S., Pastötter, B., Bäuml, K.-H., Gruber, S., Wimber, M., Klimesch, W., 2008. The electrophysiological dynamics of interference during the Stroop task. J. Cogn. Neurosci. 20, 215–25. https://doi.org/10.1162/jocn.2008.20020.

Heidlmayr, K., Kihlstedt, M., Isel, F., 2020. A review on the electroencephalography markers of Stroop executive control processes. Brain Cogn. 146. https://doi.org/10.1016/j.bandc.2020.105637

Himberg, J., Hyvärinen, A., Esposito, F., 2004. Validating the independent components of neuroimaging time series via clustering and visualization. Neuroimage 22, 1214–1222. https://doi.org/10.1016/j.neuroimage.2004.03.027

Iannaccone, R., Hauser, T.U., Staempfli, P., Walitza, S., Brandeis, D., Brem, S., 2015. Conflict monitoring and error processing: New insights from simultaneous EEG-fMRI. Neuroimage 105, 395–407. https://doi.org/10.1016/j.neuroimage.2014.10.028

Jenkinson, M., Beckmann, C.F., Behrens, T.E.J., Woolrich, M.W., Smith, S.M., 2012. FSL. Neuroimage 62, 782–790. https://doi.org/10.1016/j.neuroimage.2011.09.015

Kelly, R.E., Alexopoulos, G.S., Wang, Z., Gunning, F.M., Murphy, C.F., Morimoto, S.S., Kanellopoulos, D., Jia, Z., Lim, K.O., Hoptman, M.J., 2010. Visual inspection of independent components: Defining a procedure for artifact removal from fMRI data. J. Neurosci. Methods 189, 233– 245. https://doi.org/10.1016/j.jneumeth.2010.03.028

Lee, T.W., Girolami, M., Sejnowski, T.J., 1999. Independent component analysis using an extended infomax algorithm for mixed subgaussian and supergaussian sources. Neural Comput. 11, 417–41.

Lewis, L.D., Setsompop, K., Rosen, B.R., Polimeni, J.R., 2018. Stimulus-dependent hemodynamic response timing across the human subcortical-cortical visual pathway identified through high spatiotemporal resolution 7T fMRI. Neuroimage 181, 279–291. https://doi.org/10.1016/j.neuroimage.2018.06.056

Li, C.S.R., Yan, P., Bergquist, K.L., Sinha, R., 2007. Greater activation of the “default” brain regions predicts stop signal errors. Neuroimage 38, 640–648. https://doi.org/10.1016/j.neuroimage.2007.07.021

Li, Y.-O., Adali, T., Calhoun, V.D., 2007. Estimating the number of independent components for functional magnetic resonance imaging data. Hum. Brain Mapp. p28, 1251–66. https://doi.org/10.1002/hbm.20359

Logan, G.D., Zbrodoff, N.J., 1979. When it helps to be misled: Facilitative effects of increasing the frequency of conflicting stimuli in a Stroop-like task. Mem. Cognit. 7, 166–174. https://doi.org/10.3758/BF03197535

Logothetis, N.K., Pauls, J., Augath, M., Trinath, T., Oeltermann, A., 2001. Neurophysiological investigation of the basis of the fMRI signal. Nature 412, 150–157. https://doi.org/10.1038/35084005

Mullinger, K.J., Morgan, P.S., Bowtell, R.W., 2008. Improved artifact correction for combined electroencephalography/functional MRI by means of synchronization and use of vectorcardiogram recordings. J. Magn. Reson. Imaging 27, 607–616. https://doi.org/10.1002/jmri.21277

Mullinger, K.J., Yan, W.X., Bowtell, R., 2011. Reducing the gradient artefact in simultaneous EEG-fMRI by adjusting the subject’s axial position. Neuroimage 54, 1942–1950. https://doi.org/10.1016/j.neuroimage.2010.09.079

Mumford, J.A., Bissett, P.G., Jones, H.M., Shim, S., Ali, J., Rios, H., Poldrack, R.A., 2023. The response time paradox in functional magnetic resonance imaging analyses. bioRxiv 2023.02.15.528677. https://doi.org/10.1101/2023.02.15.528677

Nachev, P., Kennard, C., Husain, M., 2008. Functional role of the supplementary and pre-supplementary motor areas. Nat. Rev. Neurosci. 9, 856–69. https://doi.org/10.1038/nrn2478

Nigbur, R., Ivanova, G., Stürmer, B., 2011. Theta power as a marker for cognitive interference. Clin. Neurophysiol. 122, 2185–94. https://doi.org/10.1016/j.clinph.2011.03.030

Nolan, H., Whelan, R., Reilly, R.B., 2010. FASTER: Fully Automated Statistical Thresholding for EEG artifact Rejection. J. Neurosci. Methods 192, 152–162. https://doi.org/10.1016/j.jneumeth.2010.07.015

O’Connell, R.G., Dockree, P.M., Kelly, S.P., 2012. A supramodal accumulation-to-bound signal that determines perceptual decisions in humans. Nat. Neurosci. 15, 1729–1735. https://doi.org/10.1038/nn.3248

Oldfield, R.C., 1971. The assessment and analysis of handedness: The Edinburgh inventory. Neuropsychologia 9, 97–113. https://doi.org/10.1016/0028-3932(71)90067-4

Power, J.D., Mitra, A., Laumann, T.O., Snyder, A.Z., Schlaggar, B.L., Petersen, S.E., 2014. Methods to detect, characterize, and remove motion artifact in resting state fMRI. Neuroimage 84, 320– 341. https://doi.org/10.1016/J.NEUROIMAGE.2013.08.048

Power, J.D., Plitt, M., Kundu, P., Bandettini, P.A., Martin, A., 2017. Temporal interpolation alters motion in fMRI scans: Magnitudes and consequences for artifact detection. PLoS One 12, e0182939. https://doi.org/10.1371/journal.pone.0182939

Prestel, M., Steinfath, T.P., Tremmel, M., Stark, R., Ott, U., 2018. fMRI BOLD Correlates of EEG Independent Components: Spatial Correspondence With the Default Mode Network. Front. Hum. Neurosci. 12, 478. https://doi.org/10.3389/fnhum.2018.00478

Rogge, J., Jocham, G., Ullsperger, M., 2022. Motor cortical signals reflecting decision making and action preparation. Neuroimage 263, 119667. https://doi.org/10.1016/j.neuroimage.2022.119667

Scheeringa, R., Bastiaansen, M.C.M., Petersson, K.M., Oostenveld, R., Norris, D.G., Hagoort, P., 2008. Frontal theta EEG activity correlates negatively with the default mode network in resting state. Int. J. Psychophysiol. 67, 242–251. https://doi.org/10.1016/j.ijpsycho.2007.05.017

Scheeringa, R., Fries, P., Petersson, K.-M., Oostenveld, R., Grothe, I., Norris, D.G., Hagoort, P., Bastiaansen, M.C.M., 2011. Neuronal Dynamics Underlying High- and Low-Frequency EEG Oscillations Contribute Independently to the Human BOLD Signal. Neuron 69, 572–583. https://doi.org/10.1016/J.NEURON.2010.11.044

Scheeringa, R., Koopmans, P.J., van Mourik, T., Jensen, O., Norris, D.G., 2016. The relationship between oscillatory EEG activity and the laminar-specific BOLD signal. Proc. Natl. Acad. Sci. U. S. A. 113, 6761–6. https://doi.org/10.1073/pnas.1522577113

Scheeringa, R., Petersson, K.M., Oostenveld, R., Norris, D.G., Hagoort, P., Bastiaansen, M.C.M., 2009. Trial-by-trial coupling between EEG and BOLD identifies networks related to alpha and theta EEG power increases during working memory maintenance. Neuroimage 44, 1224–1238. https://doi.org/10.1016/j.neuroimage.2008.08.041

Scherbaum, S., Dshemuchadse, M., 2013. Higher response time increases theta energy, conflict increases response time. Clin. Neurophysiol. https://doi.org/10.1016/j.clinph.2012.12.007

Schmidt, J.R., 2013. Questioning conflict adaptation: Proportion congruent and Gratton effects reconsidered. Psychon. Bull. Rev. 20, 615–630. https://doi.org/10.3758/s13423-012-0373-0

Simon, J.R., 1969. Reactions toward the source of stimulation. J. Exp. Psychol. 81, 174–176. https://doi.org/10.1037/h0027448

Snipes, S., Krugliakova, E., Meier, E., Huber, R., 2022. The Theta Paradox: 4-8 Hz EEG Oscillations Reflect Both Sleep Pressure and Cognitive Control. J. Neurosci. 42, 8569–8586. https://doi.org/10.1523/JNEUROSCI.1063-22.2022

Stroop, J.R., 1935. Studies of interference in serial verbal reactions. J. Exp. Psychol. 18, 643–662. https://doi.org/10.1037/h0054651

Twomey, D.M., Murphy, P.R., Kelly, S.P., O’Connell, R.G., 2015. The classic P300 encodes a build-to-threshold decision variable. Eur. J. Neurosci. 42, 1636–1643. https://doi.org/10.1111/ejn.12936

Ullsperger, M., Bylsma, L.M., Botvinick, M.M., 2005. The conflict adaptation effect: It’s not just priming. Cogn. Affect. Behav. Neurosci. 5, 467–472. https://doi.org/10.3758/CABN.5.4.467

Ullsperger, M., Danielmeier, C., Jocham, G., 2014. Neurophysiology of performance monitoring and adaptive behavior. Physiol. Rev. 94, 35–79. https://doi.org/10.1152/physrev.00041.2012

Ullsperger, M., Von Cramon, D.Y., 2001. Subprocesses of performance monitoring: A dissociation of error processing and response competition revealed by event-related fMRI and ERPs. Neuroimage 14, 1387–1401. https://doi.org/10.1006/nimg.2001.0935

van den Wildenberg, W.P.M., Ridderinkhof, K.R., Wylie, S.A., 2012. Once Bitten, Twice Shy: On the Transient Nature of Congruency Sequence Effects. Front. Psychol. 3, 264. https://doi.org/10.3389/fpsyg.2012.00264

van Driel, J., Swart, J.C., Egner, T., Ridderinkhof, K.R., Cohen, M.X., 2015. (No) time for control: Frontal theta dynamics reveal the cost of temporally guided conflict anticipation. Cogn. Affect. Behav. Neurosci. 15, 787–807. https://doi.org/10.3758/s13415-015-0367-2

Varoquaux, G., Sadaghiani, S., Pinel, P., Kleinschmidt, A., Poline, J.B., Thirion, B., 2010. A group model for stable multi-subject ICA on fMRI datasets. Neuroimage 51, 288–99. https://doi.org/10.1016/j.neuroimage.2010.02.010

Vassena, E., Deraeve, J., Alexander, W.H., 2020. Surprise, value and control in anterior cingulate cortex during speeded decision-making. Nat. Hum. Behav. 4, 412–422. https://doi.org/10.1038/s41562-019-0801-5

Weissman, D.H., Carp, J., 2013. The congruency effect in the posterior medial frontal cortex is more consistent with time on task than with response conflict. PLoS One 8, e62405. https://doi.org/10.1371/journal.pone.0062405

Yang, Q., Pourtois, G., 2022. Reduced flexibility of cognitive control: reactive, but not proactive control, underpins the congruency sequence effect. Psychol. Res. 86, 474–484. https://doi.org/10.1007/s00426-021-01505-6

Yarkoni, T., Barch, D.M., Gray, J.R., Conturo, T.E., Braver, T.S., 2009. BOLD correlates of trial-by-trial reaction time variability in gray and white matter: a multi-study fMRI analysis. PLoS One 4, e4257. https://doi.org/10.1371/journal.pone.0004257

Yeung, N., Cohen, J.D., Botvinick, M.M., 2011. Errors of interpretation and modeling: a reply to Grinband et al. Neuroimage 57, 316–9.

